# Postharvest handling induces changes in fruit DNA methylation status and is associated with alterations in fruit quality in tomato (*Solanum lycopersicum* L.)

**DOI:** 10.1101/2020.10.08.331819

**Authors:** Jiaqi Zhou, Bixuan Chen, Karin Albornoz, Diane M Beckles

**Affiliations:** Department of Plant Sciences, University of California, Davis, CA 95616, USA

**Author notes:** Departamento de Produccion Vegetal, Universidad de Concepcion, Region del BioBio, Chile (Karin Albornoz). **Corresponding author:** Diane M Beckles, Department of Plant Sciences, University of California, Davis, CA 95616, USA. Abbreviations: MSAP, methyl-sensitive amplification polymorphism; ‘FH’, fresh-harvested fruit; ‘M’, Mature green; ‘T’, Turning; ‘5M’, fruit harvested at Mature green and stored at 5°C after 14 days; ‘5T’, ‘5M’ fruit stored at 20°C until Turning; ‘20T’ and ‘12.5T’, fruit early harvested at Mature green and stored at 20°C and 12.5°C, respectively; SRT-PCR, Semiquantitative Reverse Transcription-Polymerase Chain Reaction; RT-qPCR, Quantitative real-time PCR; TA, titratable acidity; *SlDML2*, *Solanum lycopersicum* L. *DNA demethylase 2*; *RIN*, *RIPENING INHIBITOR*.

**Keywords:** tomato fruit ripening, postharvest handling, fruit quality, DNA methylation, methyl-sensitive amplification polymorphism

## Abstract

Postharvest handling of tomato (*Solanum lycopersicum* L.), specifically low-temperature storage and early harvest are used to extend shelf life, but often reduce fruit quality. Recent work suggests that DNA methylation dynamics influences fruit ripening through the demethylase *SlDML2* gene. However, the influence of postharvest handling on DNA methylation in relation to fruit quality is unclear. This work aimed to clarify these issues by analyzing DNA methylation using methyl-sensitive amplification polymorphism (MSAP), semi-quantitative transcriptional analysis of marker genes for fruit quality (*RIN*; *RIPENING INHIBITOR*) and DNA methylation (*SlDML2*; *Solanum lycopersicum* L. *DNA demethylase 2*), and, fruit biochemical quality biomarkers. Multivariate analysis of these data supported the view that DNA methylation of fruit was influenced more by postharvest handling than ripening stage, however, fruit quality was influenced mainly by ripening. Fruit chilled postharvest were distinct in their DNA methylation state and quality characteristics, which implied that these three phenomena i.e., chilling, methylation, and quality are highly connected. In addition, different postharvest handling methods modulated *SlDML2* transcript levels but had little effect on the level of *RIN* transcripts in fruit that reached the Turning stage after early harvest, and cold storage. These data collectively helped to advance our interpretation of tomato fruit ripening. In conclusion, our findings revealed that postharvest-induced variation in fruit quality is in relation to DNA methylation. Long-term this work will help better connect physiological changes in tomato fruit to events happening at the molecular level.

## 1. Introduction

Tomato (*Solanum lycopersicum* L.) is one of the world’s most popular fresh-market vegetables (Strange et al., 2000; Gould, 2013), and is also an important research model for fleshly-fruit development (Giovannoni et al., 2017). Tomato is highly perishable after harvest (Sloof et al., 1996), and determining adequate postharvest handling and storage conditions is important for extending fruit shelf-life and reducing postharvest losses (Nasrin et al., 2008), which is necessary given the complicated modern supply chain for fresh produce.

Early harvest and low-temperature storage can delay fruit ripening and extend shelf-life, but there can often be unintended consequences, such as lower fruit sensory quality, reduced consumer satisfaction, and as a result, fewer repeat purchases (Hobson, 1987; Heuvelink, 2005; Klein et al., 2010). For example, Mature green fruit ripened off-the-vine at temperatures lower than ambient will have maximal shelf-life, but they may not have a fully realized sensory profile (Majidi et al., 2014). These poor sensory attributes compared to ‘on-the-vine’ ripened tomato occur because fruit nutrient supply from the mother plant is prematurely disrupted, and there may be additional losses of flavor- and taste-associated compounds occurring during storage which collectively leads to postharvest waste (Beckles, 2012). Further, exposing tomatoes to temperature below 10°C can severely disrupt the normal ripening program, and leads to negative attributes, a phenomenon called postharvest chilling injury (Biswas et al., 2016; Albornoz et al., 2020). This physiological disorder is cumulative in its effect, as the consequences are normally presented during rewarming, and include unusual softening, poor flavor and taste, and, visual defects such as uneven ripening, pitting and decay (Albornoz et al., 2019).

Tomato quality attributes are determined by many genetic, physiological and biochemical events that occur as the fruit ripens. This leads to the characteristic and desirable changes in color, texture, flavor and taste. Measurements of fruit firmness can act as a proxy for texture and juiciness (Saladié et al., 2007), color is an indicator of ripening stage and visual quality (Stommel et al., 2005), and the ratio of sugar-to-acid contributes to an appealing tomato taste and is used as a marker of this attribute (Anthon et al., 2011; Beckles, 2012). The biochemical changes that lead to these and other events are controlled by programmed developmental pathways (Klee & Giovannoni, 2011) that are disrupted by off-the-vine ripening and low temperature storage (Biswas et al., 2016).

Tomato fruit ripening, and therefore quality is mediated in part, by upstream changes in DNA methylation (Stower, 2012). The promoter region of many ripening genes remains methylated during early fruit development until the onset of ripening (Zhong et al., 2013). In tomato, DNA demethylation is critical for fruit ripening and quality, while in contrast, postharvest chilling reverses this process and promotes methylation of many ripening genes, and it is associated with reduced quality (Zhang et al., 2016). Two genes have important roles in these observations: *SlDML2* and *RIN* which both increase in expression during tomato fruit development. *SlDML2* encodes a DNA demethylase, that activates hundreds of ripening-related genes in tomato fruit by removing the methyl-group from their promoter region (Lang et al., 2017). One important target of *SlDML2* is *RIN* (Liu et al., 2015; Lang et al., 2017). RIN is a central fruit ripening transcription factor (Ito et al., 2008; Li et al., 2011; Qin et al., 2012; Martel et al., 2011; Karlova et al., 2014; Ito et al., 2020; Li et al., 2020) that is also, powerfully regulated by the DNA methylation levels of its promoter regions (Liu et al., 2015; Lang et al., 2017). *RIN* and RIN-induced genes and transcription factors (TFs) are suppressed through hypermethylation, in pre-ripened fruit or fruit exposed to chilling (Zhang et al., 2016).

The dilemma raised here is that consumers prefer full-flavored tomato fruits (Bruhn et al., 1991), but postharvest practices designed to extend shelf-life often reduce fruit quality. The former observations provide a cornerstone for understanding postharvest resulting in loss of flavor of tomato fruit. Here, we researched if changes in fruit DNA methylation status are induced by different postharvest practices i.e. early harvest at Mature green, and low-temperature storage, and, if there is a relationship between changes in DNA methylation and fruit quality parameters. A global overview of changes in ripening-associated genome methylation was determined using Methyl-sensitive amplification polymorphism (MSAP) (Xu et al., 2000). This is a simple method that indicates changes in methylation sites across genomes (Yaish et al., 2014). In MSAP, two restriction enzymes *Hpa*II and *Msp*I that recognize the same CCGG sequence, but with differential sensitivity to methylation at the inner or outer cytosine are used. DNA methylation status is then determined by analyzing the number and sizes of generated fragments after enzyme digestion, and then comparing fragments from tissues at different developmental stages, environmental treatment etc. to indicate how these conditions influence global methylation status (Chen et al., 2019).

To sum up, our aim was to use MSAP to clarify how industry practices influence fruit global DNA methylation levels. This is a first step to extend shelf-life while helping improve fruit quality. Long-term, reducing postharvest losses and increasing market consumption could be possible if the relationships of fruit quality and DNA methylation are understood.

## 2. Materials and methods

### 2.1 Plant handling

Micro-Tom tomato seeds were obtained from the UCD Tomato Genetics Resource Center (TGRC). Seeds were soaked in 2.7% (v/v) sodium hypochlorite for 45 min, rinsed thoroughly in running water, placed into petri dishes with damp paper towels and located in a 20°C (± 2°C) room under 16/8 hour-photoperiod for one week. Routine watering was applied every other day. Seedlings were transferred into the greenhouse at UC Davis in 2018 and 2019 under the growth temperature between 25°C to 30°C. Fruit were randomly harvested from over one hundred Micro-Tom plants.

### 2.2 Fruit sampling

“Micro-Tom” fruit were visually assessed and sampled at Mature green (MG) and Turning (T) stages (Supplementary Fig. S1) (Takizawa et al., 2014). Harvested fruits were soaked in 0.25% (v/v) sodium hypochlorite and rinsed with nanopore water, wrapped with paper towels until dry. Treatment groups are described in Fig. 2. Fresh fruit were used for firmness, color and Total Soluble Solids (TSS) analysis, while the pericarp from the remaining sampled fruit were fresh frozen in liquid nitrogen and stored at −80°C for further assessment. Six randomly chosen fresh fruit were measured for firmness, color and TSS analyses as six biological replicates, and three biological replicates were used for titratable acidity. For reducing sugar, starch, gene expression and DNA methylation assessment, six individual fruits were randomly chosen from a group of twenty fruits of uniform size, and those six fruits were pooled together as one biological replicate. A minimum of three biological replicates were prepared in each assessment.

### 2.3 Quality assessment

#### Firmness

Fruit firmness was measured using a Texture Analyzer (TA. XT Plus, Texture Technologies, Scarsdale, NY) by compressing the middle of the whole fruit 3 mm. The maximum force for each measurement was recorded.

#### Fruit Color

Objective parameters of color i.e. L, a* and b* were recorded for each individual fruit using a Konica Minolta colorimeter (Chroma Meter CR-400, Konica Minolta Sensing Americas, Ramsey, NJ, USA). Readings were obtained using a 2° observer and standard illuminant C setting in a three-dimensional color space. Standard calculations for Hue [Arc tan(b/a)] were made as described by Mclellan et al., 1995.

#### Total Soluble Solids (TSS) and Reducing Sugars

Tomato TSS include carbohydrates, organic acids, proteins, fats and minerals contents. Degrees Brix (used to assess TSS content) of each fruit was evaluated, using a portable digital Brix refractometer (Hanna Instruments, Inc.). To further test sugar content in tomato fruit pericarp, reducing sugars (fructose and glucose) were measured (Miller, 1959).

#### Titratable acidity (TA)

An aliquot of the pericarp (1-2 g) from a single fruit pericarp was added to 10 mL water and incubated at 80°C for 20 min, centrifuged at 4,000 x *g* for 15 min. The supernatant was transferred to a beaker, and water added to 75 mL. The solution was equally divided into three tubes (technical replicates) and one drop of 0.5% (w/v) Phenolphthalein Indicator (La-Mar-Ka, Inc) was added to each. TA was obtained by titrating 0.05 M NaOH manually into the solution until a faint color was visible for a few seconds.

#### Starch content

Fruit tissues were treated, homogenized and gelatinized as described by Luengwilai et al., (2010). A 500 mL digestion mixture of 200 mM sodium acetate (pH 5.5), 1-unit α-amyloglucosidase and 12 units α-amylase were added into two tubes containing the homogenized starch solution, with a third serving as a non-enzyme control. All samples were incubated at 37°C overnight to fully digest starch into glucose. The 3,5-dinitrosalicylic acid (DNS) reagent was used for assaying glucose content as described by Dong et al., (2018).

### 2.4 Gene expression

#### RNA isolation

RNA was isolated from 100 mg ground power using a Trizol-based protocol (Leterrier et al., 2008). DNase treatment was applied during extraction using TURBO DNA-free™ Kit (Life Technologies, Carlsbad, CA, USA). RNA quality and integrity were assessed by microvolume spectrophotometer and 0.8% (w/v) agarose gel electrophoresis.

#### cDNA synthesis

cDNA was synthesized from 2 μg of RNA using random primers in 20 μL reaction with the High-Capacity cDNA Reverse Transcription Kit (Applied Biosystems, Foster City, CA, USA).

#### Semiquantitative RT-PCR (SRT-PCR)

The PCR program and cDNA input amount were optimized to ensure that the targeted gene (*SlDML2* and *RIN*) and *SlACT7* (used as an internal control) PCR bands intensity on the gel were within a linear range for quantitative analysis. One microliter of cDNA product was amplified with Ampli*Taq* polymerase in the corresponding buffer. Designed primers and product sizes are listed in Table S1. Reactions were carried out in the Gene Amp PCR system 9600 (Applied Biosystems, USA) with the following program: one cycle of 10 min at 95°C, 45 s at 57°C, 45 s at 72°C, and then 25 cycles of 45 s at 95°C, 45 s at the annealing temperature (optimized for each pair of primer: 57°C for *RIN* and *SlACT7*, and 52°C for *SlDML2* and *SlACT7*), 1 min at 72°C, following by 72°C extension for 2 min. PCR products were directly loaded into a 2% (w/v) agarose gel, electrophoresed for 45 min at 84 V, and stained with ethidium bromide. Two amplified bands were separated for each sample. The relative expression level was calculated by the ratio of intensity areas in agarose gel between the targeted gene and *SlACT7* by Image J (Schneider et al., 2012). All enzymes and buffers were from Applied Biosystems (USA).

#### Quantitative real-time PCR (RT-qPCR)

cDNA created as described was diluted 80-fold. RT-qPCR was performed in a 10 μL reaction using iQ™ SYBR^®^ Green Supermix (Bio-Rad, Hercules, CA, USA). Applied Biosystems 7300 Real Time PCR system (Applied Biosystems, USA) was used. Primers were designed based on the cDNA sequences (Table S2). The efficiency for all pairs of primers was close to 100%, so the comparative Ct Method (ΔΔCT Method) was applied for analyzing data. Three biological replicates and three technical replicates per bio-replicate were used for each experimental condition.

### 2.5 Methyl-sensitive amplification polymorphism (MSAP)

A schematic of the MSAP procedure is shown in Fig. 1, while detailed steps in the protocol are described below.

**Fig. 1.**
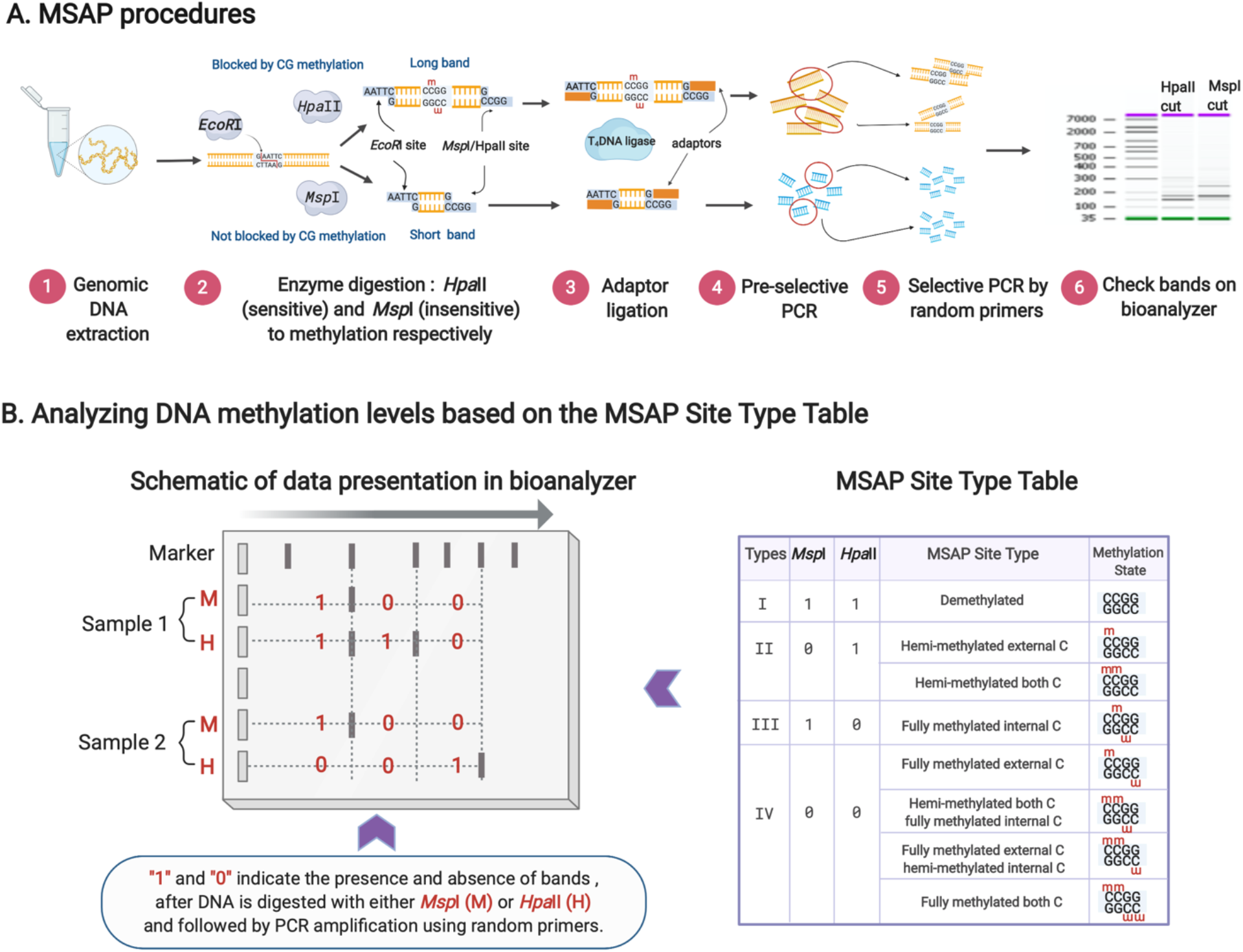
Methyl-sensitive amplification polymorphism (MSAP) procedures. (**A**) Individual steps used in the MSAP protocol. For details refer to Section 2.5 in the materials and methods. (**B**) Schematic representation of the DNA fragments and their classification as one of four ‘site-types’ determined by the *Msp*I (M) and *Hpa*II (H) digestion pattern. Each pattern represents a different methylation state shown in the MSAP Site Type Table (adapted) (Fulneček & Kovařík, 2014; Guarino et al., 2019).

#### DNA extraction

Three biological replicates were used for MSAP analysis, each derived from a pool of six individual fruits. Genomic DNA from tomato fruit pericarp was extracted by optimizing published CTAB protocols (Yan et al., 2018). Three hundred milligram of frozen powder was added to 1 mL of chilled washing buffer (100 mM Tris-HCl (pH 8.0), 5 mM ethylenediaminetetraacetic acid (EDTA, pH 8.0), 0.35 M glucose, 1% (w/v) polyvinylpyrrolidone (PVP), 2% (v/v) β-mercaptoethanol), put on ice for 10 min, and centrifuged 2 min to remove all supernatant. This step helps remove high levels of polysaccharides and polyphenols. Then 1 mL 65°C prewarmed CTAB buffer (100 mM Tris-HCl (pH 7.5), 25 mM EDTA, 1.5 M NaCl, 2% (w/v) CTAB, and 0.3% (v/v) β-mercaptoethanol) was added into each sample. The remaining steps were done as described by Healey et al., (2014). The DNA pellet was dissolved using 50 μL nuclease-free water. The absorbance ratio 260/280 nm and 260/230 nm were between 1.8 and 2.0. DNA appeared on an 0.8% (w/v) agarose gel as a single high molecular weight band.

#### Genomic DNA digestion

Two aliquots of 1 μg genomic DNA from each sample were treated by 20 U each of HF-*EcoR*I and either methylation-sensitive *Hpa*II or methylation-insensitive *Msp*I with Cutsmart buffer in a total volume of 30 μL. The reaction was held at 37°C for 4 hours, followed by heating at 80°C for 10 min to deactivate the digestion enzyme. All restriction enzymes and buffers were from New England Biolabs (USA).

#### Adaptor ligation

Two pairs of adaptors were designed for HF-*EcoR*I and *Hpa*II*/Msp*I (See Table S3). Fifty pmol of each adaptor pair was placed in a total volume of 40 μL, heated at 72°C for 10 min and all tubes were then cooled in a tightly closed box overnight. Then, 15 μL of digested DNA, annealed adaptor pairs, T_4_ ligase and ligase buffer, in a total volume of 30 μL, was incubated overnight at 18°C.

#### Preselective PCR amplification

Twenty-five μL of the ligation was added to the PCR reaction, which included the HF-*EcoR*I pre-selected primer, *Hpa*II*/Msp*I pre-selected primer, dNTPs, Ampli*Taq* DNA polymerase (Applied Biosystems, USA) and PCR buffer I, in a total volume of 30 μL. The program was comprised of one cycle of 5 min at 72°C, 3 min at 94°C, and then 36 cycles of 30 s at 94°C, 1 min at 56°C, 1 min at 72°C, and 10 min at 60°C. Nine microliters of the PCR reaction product was checked on a 1.5% (w/v) agarose gel after electrophoresis at 84 V for 45 min. A DNA ‘smear’ of even intensity was observed among all DNA samples, verifying successful digestion and ligation.

#### Selective PCR amplification

The pre-selective PCR product was diluted 10-fold and used as a template in the following PCR reaction. Each template was loaded twice into two different tubes applying two pairs of selective primers separately for generating diverse data. Each PCR reaction also included a pair of selective primer, dNTPs, Ampli*Taq* DNA polymerase (Applied Biosystems, USA) and PCR buffer I, in a total volume of 20 μL. All primer sequences were listed in Table S1. The touchdown program was set as following: 1 cycle of 45 s at 94°C, 30 s at 65°C, and 1 min at 72°C, 12 cycles with decreasing the annealing temperature by 0.7°C per cycle, and then 20 cycles of 30 s at 94°C, 30 s at 55.9°C, and 1 min at 72°C.

#### Analyzing selective PCR bands number and size

PCR products were purified using MinElute^@^ PCR purification kit (QIAGEN, USA) which collects fragments from 70 bp to 4 kb. The fragments were diluted to reach a concentration range of 1-10 ng/μL as required by Agilent High Sensitivity DNA kit. A 2100 Bioanalyzer (Agilent Scientific Instruments, USA), with a micro-capillary based electrophoretic cell, was used to analyze the PCR bands.

#### Data analysis

Three biological replicates were performed for each tomato treatment. A “1” or “0” indicates the presence and absence of a DNA band respectively. If at least two replicates of the three showed a PCR band, this was classified as a “1”, while, if only one or no replicate showed a band, this was considered as “0”. Alteration in DNA methylation of tomato fruit was determined by comparing the site type (see Fig. 1B and Table S4) from two different treatments, and the definition of “*de novo* methylation”, “demethylation” and “no change” was according to Chen et al., (2019).

### 2.6 Statistical analysis

All statistical analyses were done using the R platform (R Core Team, 2020). Box plots for fruit quality and gene expression, and the heatmap of DNA methylation patterns (Fig. 3) were generated by ggplot2 (Wickham, 2009). Compact Letter Display of Pairwise Comparisons (CLD) (Piepho, 2004) were used for testing all pairwise comparisons of least-squares mean, with the significance level set at 0.05. Hierarchical Clustering analysis (HCA) was performed using “Euclidean distance” by the function of “dist” and “hclust” (Murtagh & Legendre, 2014). Principal Component analysis (PCA) for MSAP data were created by “mixOmics” (Rohart et al., 2017) and “tidyverse” (Wickham et al., 2019), and PCA for quality parameters was completed by MetaboAnalystR (Pang et al., 2020).

## 3. Results

To explore potential changes at the molecular level, and in tomato fruit quality due to postharvest practice, specifically, early harvest and low-temperature storage, we harvested fruit at Mature green (the earliest harvest stage), and stored them at different temperatures (20°C, 12.5°C) until they reached the Turning stage (Fig. S1 and Fig. 2). Turning is the ripening stage just before Red ripe for ‘Micro-Tom’, which is similar to the Pink stage for conventional tomatoes (United States Department of Argiculture, 1975). To investigate postharvest chilling injury, we stored Mature green fruit at 5°C for 14 days in order to induce this disorder (Albornoz et al., 2019). Tomato fruit are unable to ripen at this temperature, so after chilling they were allowed to recover at 20°C until they reached Turning. By comparing postharvest fruit ripened under different temperatures to those ripened ‘on-the-vine’, postharvest handling induced alterations might be uncovered.

**Fig. 2.**
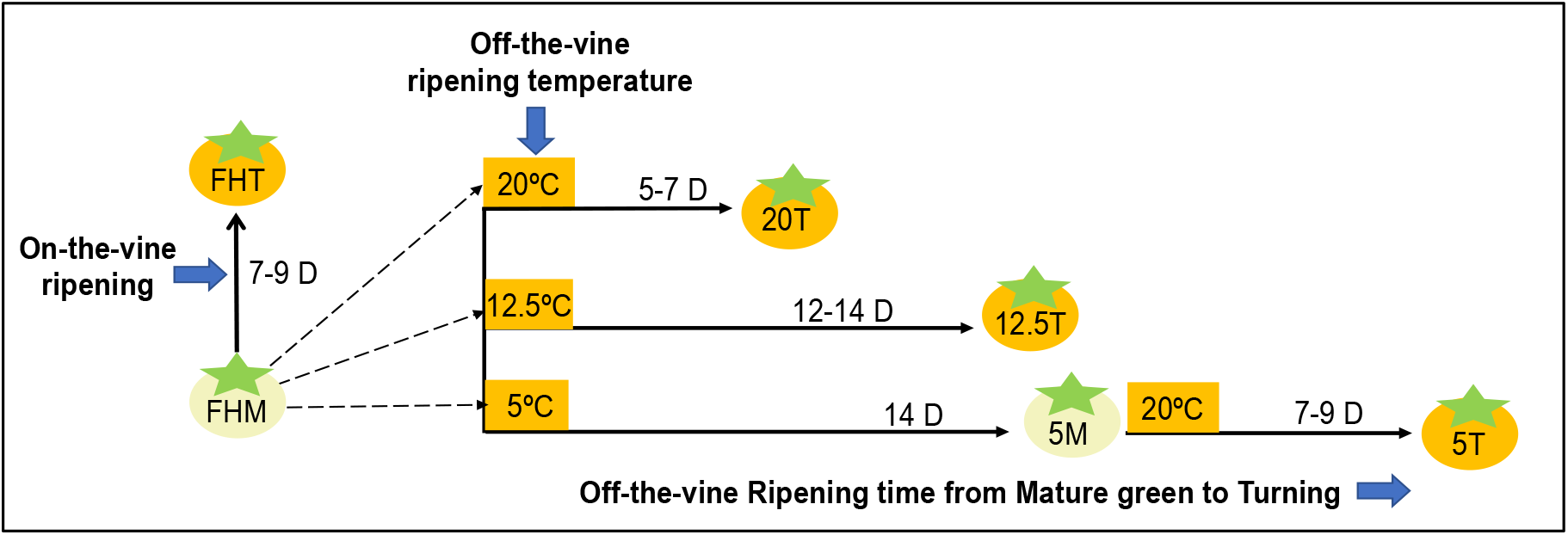
A diagram to illustrate the different fruit treatments used in this study. Different comparisons were made based on the following: **1) Fruit development:** Tomato fruit were freshly harvested at Mature green (M) and Turning (T) stages and are described as ‘FHM’ and ‘FHT’, respectively. **2) Temperature treatment**: Fruit were harvested at M, stored at 20°C or 12.5°C, and sampled until Turning. These fruit are described as ‘20T’ and ‘12.5T’, correspondingly. **3) Chilling injury**: ‘5M’ represents fruit that were harvested at M, stored at 5°C for 14 days, and directly sampled. These ‘5M’ fruit were rewarmed at 20°C until Turning and described as ‘5T’. **4) Ripening time**: The numbers beside each line indicate the timeframe between samples.

### 3.1 DNA methylation variation due to ripening and postharvest handling

The DNA methylation data generated by MSAP were grouped into one of three classes based on treatment-induced changes i.e., *de novo* methylation, demethylation, or no change in methylation status. These data were depicted in two ways; first, showing details of the relative abundance of individual sites (DNA bands) that led to the above classification (Figure 3), and second, providing an overview of the data i.e. the percentage of each methylation class induced by the treatment (Figure 4).

**Fig. 3.**
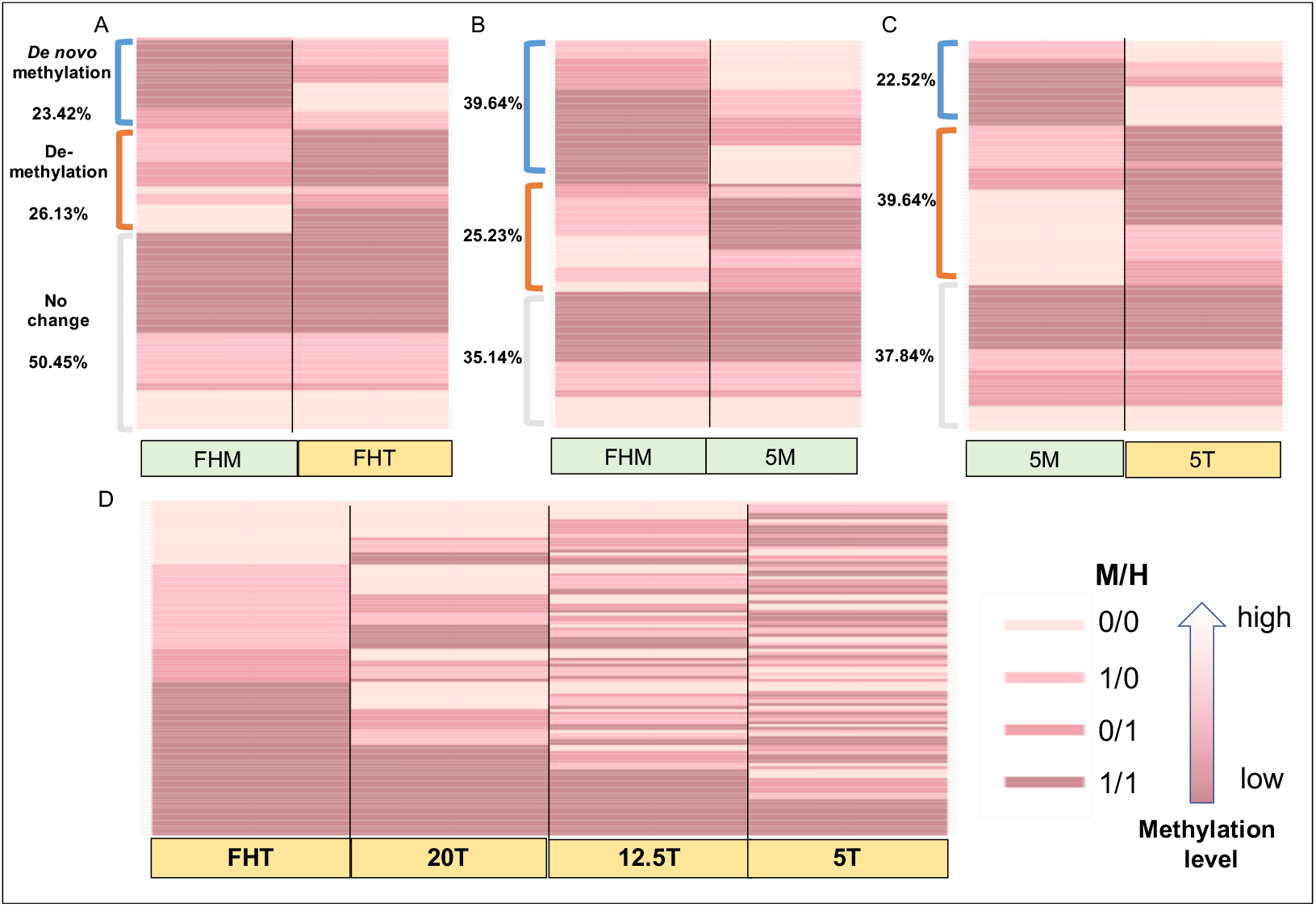
Global DNA methylation status as determined by the banding pattern generated by MSAP. Bands are shown as one of four colors from dark to light pink indicating low to high DNA methylation levels respectively. Each color indicates a distinct DNA methylation state, illustrated in the MSAP site type table (Fig. 1B and Table S4). Each row represents one band of a specific size generated by restriction with *Msp*I (M) or *Hpa*II (H). A total of 111 bands (rows) were generated by the MSAP. All figures show the same 111 bands but arranged differently for ease of comparison. In Figs. 3A, 3B and 3C, bands were ordered and organized based on the three DNA methylation patterns observed when the two samples were compared i.e. ‘No change’ in methylation status (grey bracket), ‘Demethylation’ (orange bracket) and ‘*de novo* methylation’ (blue bracket). For Fig. 3D, bands were organized by their similarity to ‘FHT’. Comparisons examined were: **(A) Normal ripening**: comparison between fresh-harvest Mature green fruit ‘FHM’ and fresh-harvest fruit at Turning i.e. ‘FHT’. **(B) Chilling effect**: fresh-harvest Mature green fruit ‘FHM’ compared to Mature green fruit stored at 5°C i.e. ‘5M’. **(C) Rewarming effect**: when ‘5M’ fruit were stored at 20°C until Turning i.e. ‘5T’. **(D) Ripening and Treatment**: all samples at Turning stage i.e. ‘FHT’, ‘20T’, ‘12.5T’ and ‘5T’, were compared.

**Fig. 4.**
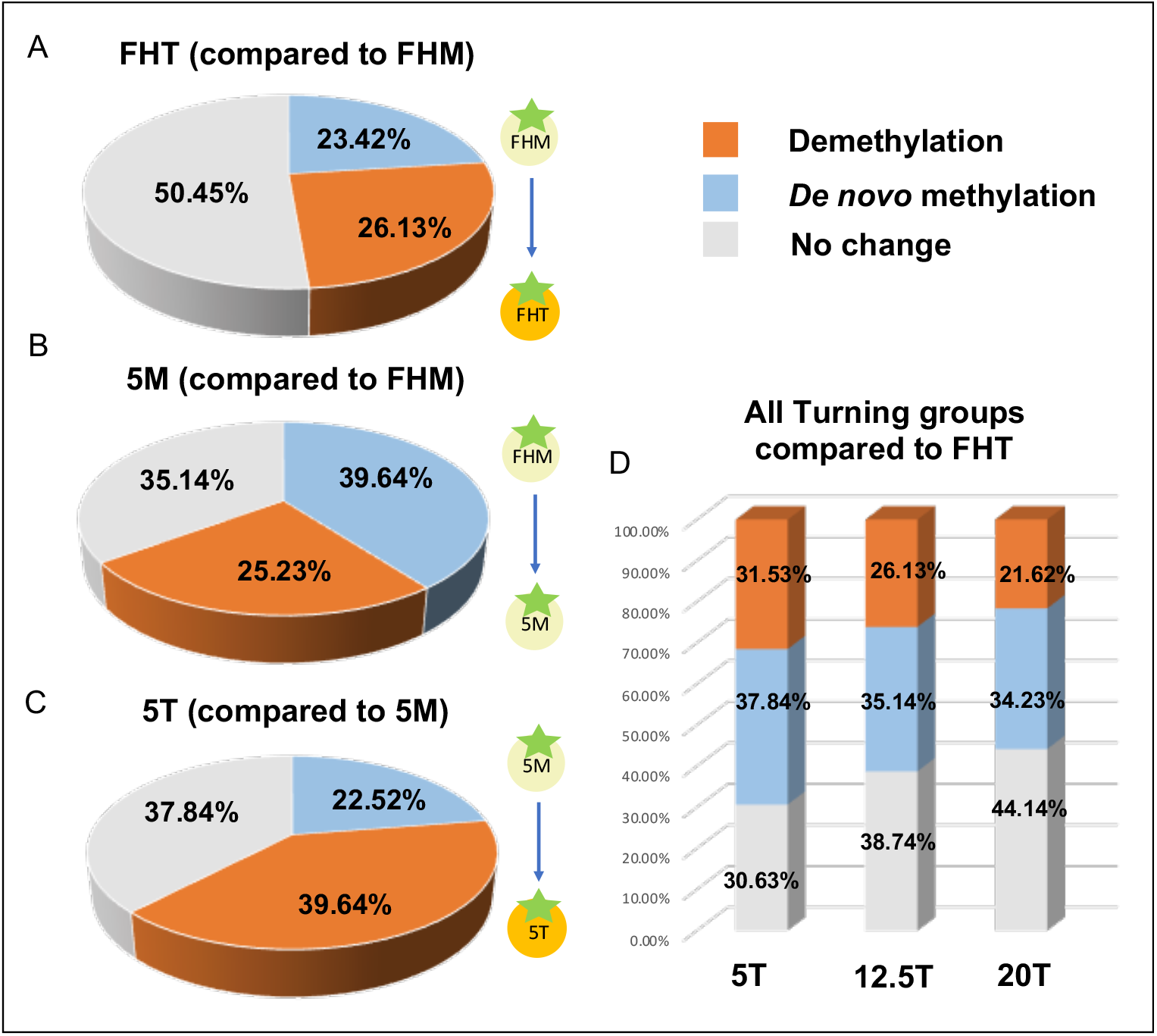
Comparisons of fruit DNA methylation status due to ripening and chilling. **The following factors were examined: (A) Normal ripening**: fruits harvested at T (‘FHT’) were compared to fruits at M (‘FHM’). **(B) Chilling**: fruits at Mature green stored at 5°C after 14 days (‘5M’) were compared to fruit before they were chilled (‘FHM’). **(C) Rewarming**: chilled fruit rewarmed at 20°C until T (‘5T’) were compared to chilled fruits (‘5M’). **(D) Postharvest ripening**. Comparisons were made between fruit harvested at Mature green and allowed to ripen under different conditions until Turning (‘5T’, ‘12.5T’, ‘20T’) compared to ‘FHT’.

We found distinctive global DNA methylation levels in fruit at the same ripening stage i.e. Turning, due to postharvest handling. This conclusion was drawn based on the MSAP data (Fig. 3D and Fig. 4D), which indicated that there were different *de novo* methylation and demethylation events occurring across assorted Turning fruit, and only a limited number of the examined sites did not change. ‘FHT’ fruit were ripened under optimal condition. Early-harvested fruit that were ripened at 20°C (‘20T’) was the most similar in methylation patterns to ‘FHT’, while the most contrastable DNA methylation fragmentation atlas was observed between ‘FHT’ and ‘5T’.

Comparing the methylation levels in ‘FHM’ vs. ‘FHT’, we found that 26.13% of the genomic sites tested represented DNA demethylation events, which is greater than the percentage representing *de novo* methylation events (23.42%) (Fig. 3A and Fig. 4A). This indicates that the overall cytosine methylation was reduced during ‘on-the-vine’ ripening.

To focus on the chilling effect on Mature green fruit, chilled ‘5M’ were compared to those that were freshly harvested ‘FHM’ (Fig. 3B and Fig. 4B). There were 39.64% *de novo* methylation events found in ‘5M’. This value was higher than the percentage of sites which were unchanged (35.14%), and even higher than those that were demethylated (25.23%). Thus, we confirmed that chilling led to markedly higher DNA methylation level.

After transferring previously chilled fruit to room temperature, fruit were able to resume ripening. Under this recovery treatment, more genomic sites (39.64%) were demethylated, compared to those that underwent *de novo* methylation (Fig. 3C and Fig. 4C). The high percentage of demethylation events during rewarming (from ‘5M’ to ‘5T’) is similar to the trend seen during normal ripening (from ‘FHM’ to ‘FHT’). We also observed that DNA methylation was reversible when fruit were rewarmed after chilling, since *de novo* methylation events increased in ‘5M’, but after rewarming until Turning, demethylation occurred.

To summarize all observations from the MSAP analysis, (a) postharvest handling induced changes in fruit global DNA methylation status, and the rank of their methylation status based on similarity to “on-the-vine” ripened fruit at Turning was ‘20T’ > ‘12.5T’ > ‘5T’; (b) demethylation occurred as the fruit ripened; (c) chilling at the onset of fruit ripening resulted in hypermethylation and the inhibition of fruit ripening; and (d) DNA demethylation occurred in the following rewarming process, but some of these demethylated sites were different to the fruit under normal ripening.

### 3.2 Tomato fruit quality is largely influenced by early harvest and low-temperature storage

We examined the quality of the fruit harvested and stored, using common postharvest markers i.e. firmness, total soluble solids (TSS), reducing sugars, objective color, titratable acidity, and starch content. We observed that ‘FHT’ and ‘20T’ showed optimal quality, while ‘5T’ was comparatively the worst compared with other Turning fruit (Fig. 5). The group of fruit stored at 12.5°C (‘12.5T’), was better than ‘5T’ but lower than ‘20T’ based on the measured quality parameters.

**Fig. 5.**
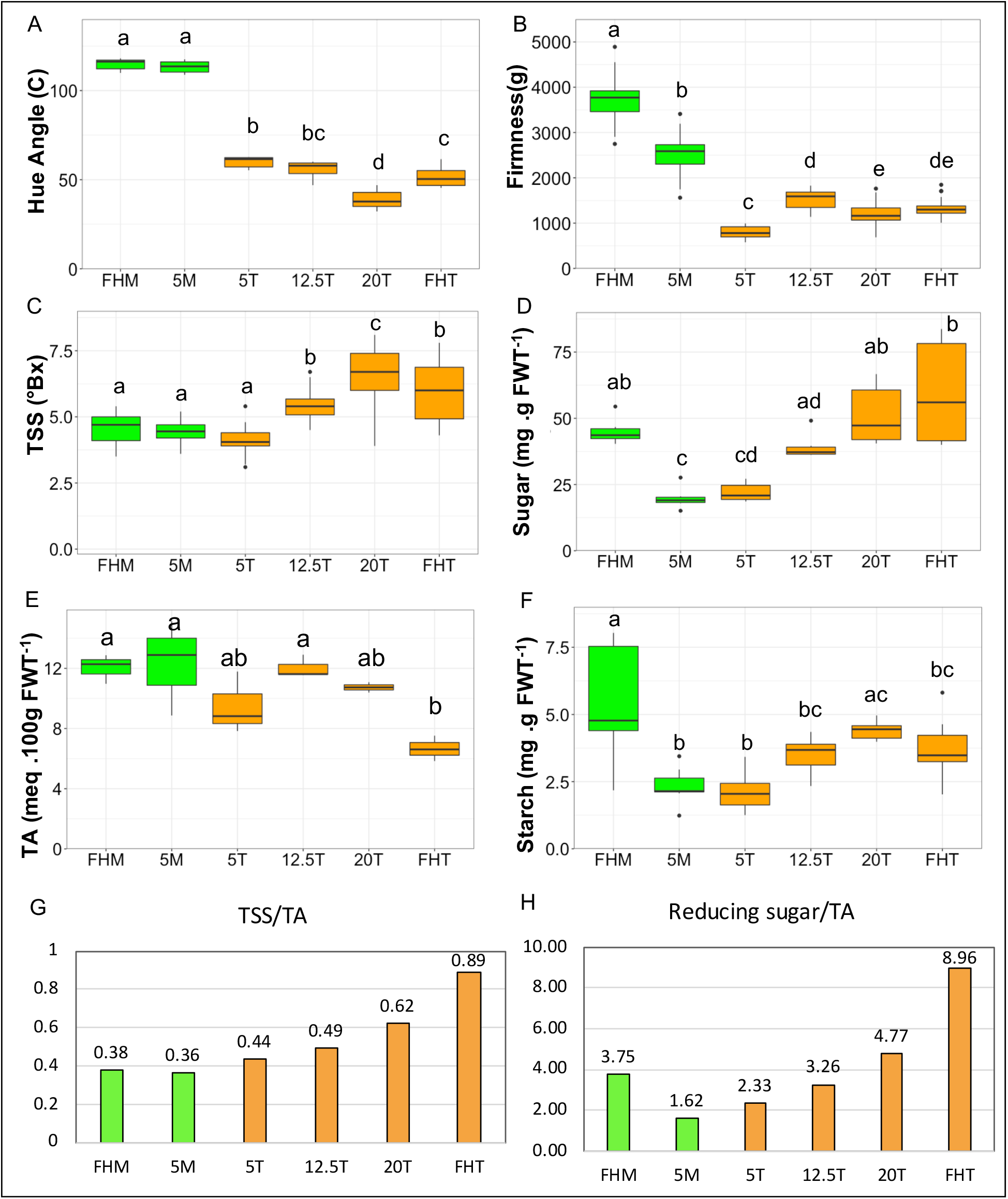
Fruit quality assessment. Six groups, including **‘** FHM’, ‘5M’, ‘5T’, ‘12.5T’, ‘20T’ and ‘FHT’, were analyzed. Letters above the box indicate significant differences across all groups (*p* < 0.05, CLD). **(A)** Objective color: Hue angle (°). **(B)** Firmness (g). **(C)** Total soluble solids (TSS) (°Bx). **(D)** Reducing sugars, represented by mg. sugar g^−1^ (fresh weight). **(E**) Titratable acidity (TA), represented by meq. 100g^−1^ (fresh weight). **(F)** Starch content, represented by mg. starch g^−1^ (fresh weight). **(G)** TSS to TA ratio, was calculated using the mean value of each treatment. **(H)** Reducing sugars to TA ratio, was calculated by the mean value of each treatment.

#### Time to ripen

Low-temperature storage slows down fruit ripening, while chilling temperatures i.e. < 10°C arrest the process (Gonzalez et al., 2015). In this work (Fig. 2), fruit harvested at Mature green and stored at optimum ripening temperature 20°C (‘20T’) took on average 5 to 7 days to reach Turning, which is faster than ‘on-the-vine’ ripening (7-9 days). Tomato chilled at 5°C for 2 weeks reached Turning when stored at 20°C, taking 7 to 9 days on average. Remarkably, non-chilling, but low-temperature storage resulted in delayed ripening in ‘12.5T’. It took around 12 to 14 days to reach Turning, which is the longest timespan of all groups.

#### Color and firmness

Fruit ripening stages are often defined by external color (Gonzalez et al., 2015), e.g., in our study, all fruit were categorized as reaching ‘Turning’ when they attained the established predetermined color that defines that stage (Fig. S1). Objective color (hue angle) was used to quantify the green to red hues characteristic of ripening; hue angle is low in red, and high in green fruit respectively. In Fig. 5A, hue angle was identical in the green ‘FHM’ and ‘5M’ fruit, and higher than all fruit at Turning. Notably, the hue angle of ‘20T’ fruit was lower (redder), compared to all other Turning fruit including ‘FHT’. Ripening and carotenoid accumulation may have been accelerated in harvested fruit stored at 20°C in a controlled environment (see Fig. 2; Suslow & Cantwell, 2002). ‘FHT’ ripened in the greenhouse would have experienced variability in ambient light and temperature, which may have retarded carotenoid accumulation (Gautier et al., 2008), relative to ‘20T’. It should be noted that although fruit color was used here as a marker for fruit ripening stage and quality, other parameters that are also important for quality may not change in concert with color, especially after interference of the ripening program by postharvest treatments (Deltsidis et al., 2018; Shewfelt et al., 1988; Lana et al., 2005)

All Mature green fruit were firmer than those at Turning (Fig. 5B). ‘FHM’ fruit were firmer than ‘5M’ fruit, while ‘5T’ fruit was softest of all the fruit examined. The abnormal changes in fruit firmness were caused by chilling in ‘5M’ (Biswas et al., 2016), and the excessive softening in ‘5T’ fruit is a classic symptom of postharvest chilling injury which appears after the transfer of produce from low to warm temperatures (Cheng & Shewfelt, 1988).

#### Total Soluble Solids (TSS), Sugar and titratable acidity (TA)

Tomato fruit at Turning have increased TSS, sugar and decreased starch content compared to the onset of ripening (Luengwilai & Beckles 2009; Luengwilai et al., 2010). Our data were in accordance: ‘on-the-vine’ ripening was associated with higher TSS and higher reducing sugar when ‘FHT’ fruit was compared to ‘FHM’ (Figs. 5C and 5D). However, TSS content increased marginally from 4.56% in ‘FHM’ to 5.93% in ‘FHT’, which may be due to the high acid content in Micro-Tom which contributes to TSS (Luengwilai et al., 2010).

The sugar-to-acid ratio influences fruit taste, and is also a fruit maturity indicator, i.e. lower ratio indicates retarded maturity (Tigist et al., 2013; Beckles, 2012). ‘5M’ and ‘5T’ had significantly lower sugar compared to ‘FHM’, indicating that chilling and early harvest results in radical sugar consumption (Fig. 5D). Interestingly, after rewarming, the sugar content of ‘5T’ was similar to that in ‘5M’ fruit, which means that chilling resulted in a depletion of sugars that could not be restored when the fruit were allowed to ripen at room temperature. This observation is consistent with the lower glucose and fructose content in ‘Micro-Tom’ fruit induced by chilling (Gómez et al., 2009).

TA content from ‘FHM’ to ‘FHT’ decreased significantly (*p* < 0.05), i.e. from 12.0 and 6.7 meq. 100g^−1^ FW (see Fig. 5E), which is in agreement with other work (Teka, 2013). However, decreased acidity, a normal feature of tomato fruit ripening did not occur in any other fruit samples i.e. ‘5T’, ‘12.5T’ and ‘20T’, indicating poorer quality induced by the postharvest handling. The TSS-to-TA ratio and sugar-to-TA ratio (Figs. 5G and 5H) illustrate the rank of fruit taste as ‘FHT’ > ‘20T’ > ‘12.5’ > ‘5T’.

#### Starch

Green fruit store high levels of starch, which is degraded to sugars during ripening. When fruit are harvested at Mature green, starch becomes the primary carbon and energy source for ‘off-the-vine’ fruit development (Beckles, 2012). The data shown in Fig. 5F followed what was expected, but remarkably, starch content at ‘5T’ and ‘5M’ were identical. Thus, rewarming previously chilled fruit didn’t influence starch and reducing sugar content significantly, which is unlike the trends in ‘12.5T’, ‘20T’ and ‘FHT’ fruit. This phenomenon was also seen in Albornoz et al., (2019), where starch degradation occurred during cold-storage fruit, but there was no resumption during the rewarming period. In many plant tissues, chilling accelerates starch degradation presumably as a response to stress (Dong & Beckles, 2019).

### 3.3 Transcriptional analysis of DNA demethylase *SlDML2* and the master ripening regulator *RIN* in fruit ripened under different conditions

Examining the transcriptional levels of *SlDML2* may contribute to an understanding of DNA methylation dynamics due to postharvest handling. The *SlDML2* transcriptional activity was generated by Semi-quantitative RT-PCR (SRT-PCR) and Quantitative Real-Time PCR (RT-qPCR). SRT-PCR can verifiably and efficiently allow for qualitative comparisons of transcripts with relatively high accuracy and at lower costs than RT-qPCR with the necessary optimization (Marone et al., 2001; Romero et al., 2007; Antiabong et al., 2016). The data generated from SRT-PCR was largely in agreement with that from RT-qPCR (Fig. S2).

RIN is a key controller of tomato fruit ripening and quality by regulating the expression of some, but not all ripening genes connected to climacteric ethylene production (Ito et al., 2020; Li et al., 2020). We hypothesized that assessing *RIN* transcriptional levels will help to understand the effect of postharvest handling on fruit ripening process. As shown in Fig. 6B, *RIN* transcripts were not detected in Mature green fruit, but were high, and identical among the different Turning groups. We also found no *SlDML2* transcript in Mature green fruit, only in Turning fruit (see Fig. 6A). Surprisingly, the highest transcript levels of *SlDML2* were in ‘12.5T’ fruit, even higher than ‘20T’ and ‘FHT’.

**Fig. 6.**
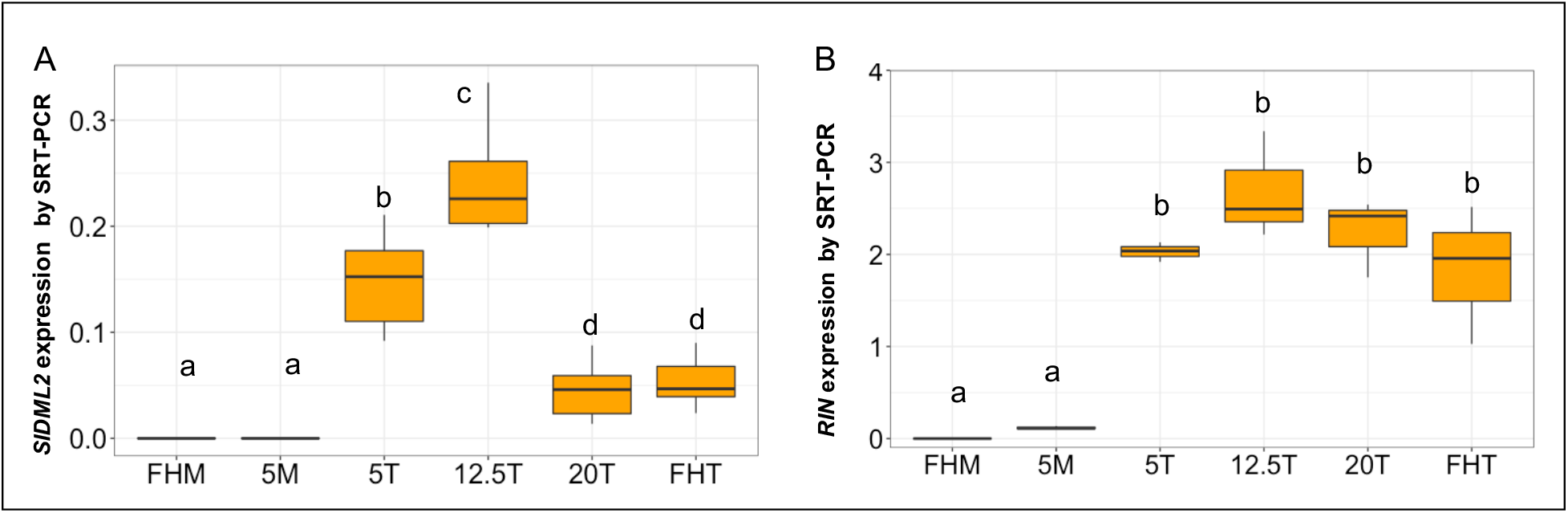
Gene expression levels of *SlDML2* and *RIN* by Semi-quantitative RT-PCR. Three biological replicates were included, in which *SlACT7* was the reference gene. Letters above the box indicate significant differences across all groups (*p* < 0.05, CLD). (**A**) *SlDML2* relative expression. (**B**) *RIN* relative expression.

Analysis of the fruitENCODE database (www.epigenome.cuhk.edu.hk/encode.html) (Lü et al., 2018), indicates that *RIN* is expressed after Mature green, increases and reaches maximal levels at Pink (equivalent to the Turning stage in ‘Micro-Tom’) (Fig. S4). Our data was identical, i.e. *RIN* expression was low in green fruit, but high in all Turning fruit, regardless of postharvest treatments. This suggests that *RIN* transcriptional levels may be determined more by final ripening stage rather than by the preceding postharvest storage conditions.

In ‘Ailsa Craig’ fruit, *SlDML2* expression was low relative to *RIN*, but it increased almost 3-fold at the onset of Mature green, increased further during Breaker, and decreased thereafter (Fig. S3). Previous work showed that *SlDML2* increases from immature to Breaker stage, peaks at Turning, and decreases at Red ripe in a cherry tomato (Liu et al., 2015). In our current study, Turning fruit ripened at 12.5°C had the highest transcript levels of *SlDML2* along with the long timespan (12-14 days) in the transition from Mature green to Turning stage. Taken together, it can be deduced that during this long-period of low-temperature storage, *SlDML2* transcripts progressively and gradually increased, which is required for the fruit to ripen before the expression peak that usually occurs at Turning. However, the *SlDML2* transcripts of other Turning fruit (‘FHT’, ‘20T’ and ‘5T’) are low, because expression was less nuanced due to accelerated ripening. This indicates that chronological factors were important for the expression of this gene.

### 3.4 Multivariate analysis according to DNA methylation and fruit quality

To find patterns that could indicate connections between DNA methylation and fruit quality among the groups of examined tissues, we performed multivariate analyses of the MSAP data using hierarchical clustering analysis (HCA) and principal component analysis (PCA). In the MSAP HCA plot (Fig. 7A), ‘5T’ and ‘5M’ fruit clustered as one group; these fruit interestingly were both chilled, and both showed symptoms of postharvest chilling injury with abnormal firmness (Fig. 5B). The ‘12.5T’ and ‘20T’ fruit clustered into another group, and they were ripened postharvest at non-chilling temperatures. The ‘FHT’ and ‘FHM’ clustered as a distinct group, and they were both ‘fresh-harvested’. HCA of fruit quality was starkly different to that of the MSAP data. HCA presented the fruit samples as two clusters that are separated only by ripening stage (Fig. 7B). However, there was some separation among Turning fruit based on ripening temperature.

**Fig. 7.**
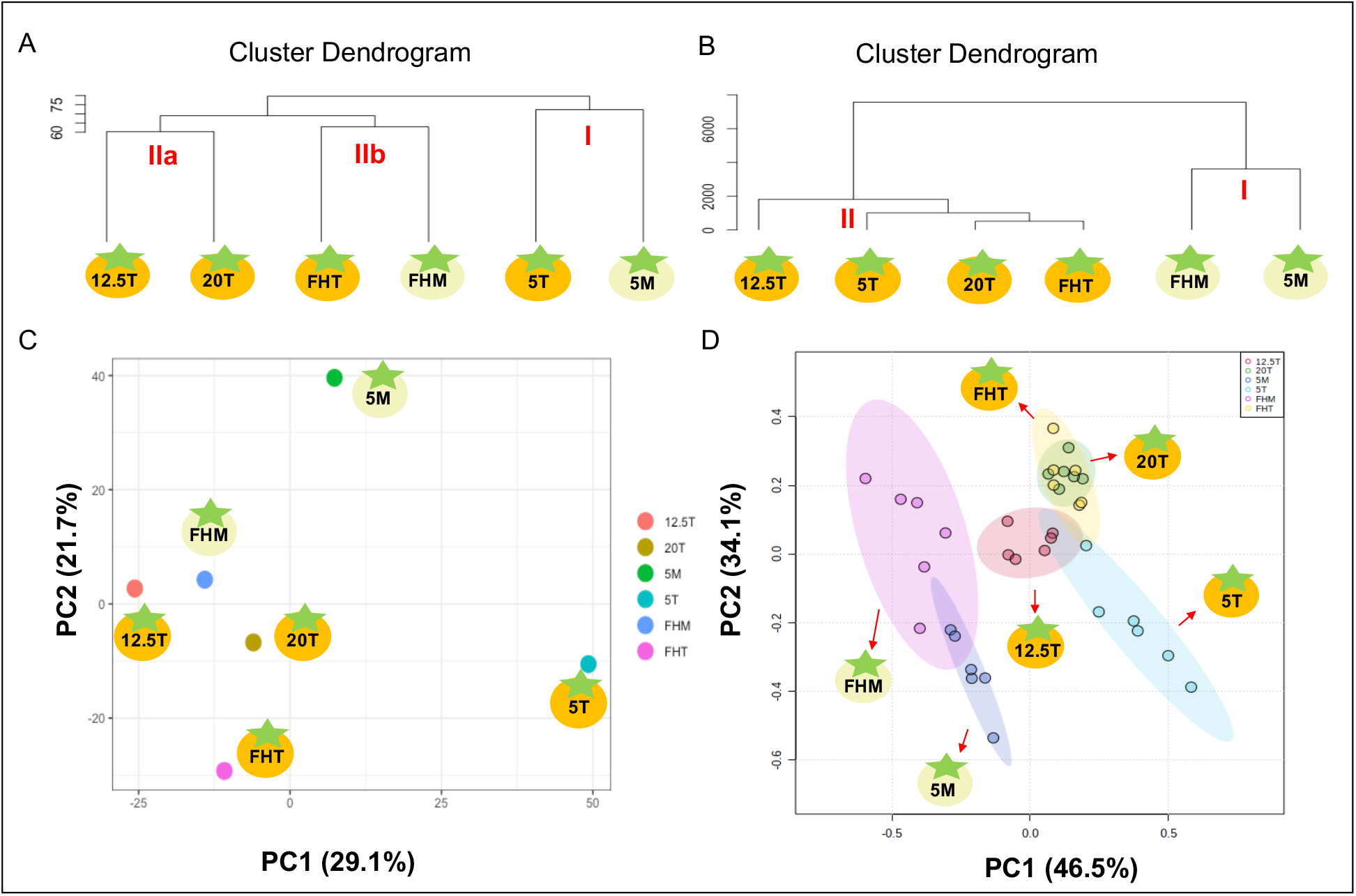
**Hierarchical clustering analysis (HCA) of (A) DNA methylation and (B) fruit quality.** The calculation for clustering was based on a matrix of site types of (A) methylation status and (B) quality parameters, including color, firmness, TSS, reducing sugar, starch, and TA. Clusters I, and II are labeled in red. **Principal component analysis (PCA) of (C) DNA methylation and (D) fruit quality.** The distance in the PCA plots shows their relationship among the variables. The calculation for clustering was based on the same matrix used in (A) and (B), and the top two PCs, were chosen to generate the 2-D plot.

The PCA of the MSAP data, showed less nuance than the HCA data – only ‘5M’ and ‘5T’ were distinct and clustered away from other groups, while ‘FHM’ was closer to the other Turning groups. PCA of tomato fruit quality (Fig. 7D), matched the HCA results i.e. green fruit ‘5M’ and ‘FHM’ formed a distinct cluster away from the Turning fruit. Still on the fruit-quality PCA, ‘5T’ showed some distinction from the other Turning fruit.

## 4. Discussion

DNA methylation has been reported to play a critical role in regulating fruit ripening. In the current work, we focused on potential changes in DNA methylation dynamics due to postharvest handling. Many postharvest strategies are designed to extend fruit shelf life, but often result in loss of fruit quality, and may unintentionally contribute to postharvest waste. While the relationship between DNA methylation and tomato fruit ripening is well understood (Shinozaki et al., 2018), it was not known what effect early harvesting and storage temperature, which often disrupt the ripening program, would have on DNA methylation and the expression of the key genes in this process. Our work demonstrated that early harvest and postharvest storage temperatures greatly influenced the speed of fruit ripening, fruit quality and DNA methylation levels, but that the relationship was not linear.

During tomato fruit development and especially during the transition from green fruit to red that occurs during ripening, many genomic demethylation events trigger the expression of ripening-related genes (Giovannoni et al., 2017). We found more demethylation events (26.13%) in Turning fruit compared to Mature green fruit, consistent with data from cv. Alisa Craig (Teyssier et al., 2008; Zhong et al., 2013). This suggests that ripening-induced DNA demethylation is conserved in tomato cvs. ‘Micro-Tom’ and ‘Alisa Craig’, even though ‘Micro-Tom’ has comparatively more methylated regions (Lang et al., 2017).

The chilling-induced hypermethylation detected in this study, explains why postharvest chilling inhibits ripening in Mature green fruit (Biswas et al., 2016). Similar to chilled red fruit (Zhang et al., 2016), chilling green fruit induced hypermethylation, presumably of the promoter regions of many ripening-related genes. The expression of these genes may be regulated by RIN, but methylation also inhibits RIN’s actions, and would delay ripening. Rewarming the chilled Mature green fruit gave rise to DNA demethylation, which is a prerequisite for ripening (Lang et al., 2017). However, not all sites methylated due to chilling were reversible after rewarming, and explains why ‘5T’ fruit were of poorer quality than other Turning fruit. This is in agreement with the growth rate and methylation pattern of chilled and rewarmed cucumber radicles (Chen et al., 2019).

By incorporating all measurements, the rank in fruit quality at Turning from best to poorest, was: ‘FHT’ > ‘20T’ > ‘12.5T’ > ‘5T’ (Fig. 5), opposite to changes in cytosine methylation levels where ‘20T’ < ‘12.5T’ < ‘5T’ (Fig. 3D and Fig. 4D) when compared to ‘FHT’. Vine-ripened fruits import nutrients until harvest, while postharvest-ripened fruit are prematurely removed from their source of nutrients. Low-temperature storage further disrupts the ripening program of these harvested fruit. It may be inferred that changes in methylation events are integral to how these anthropogenic factors affect fruit biological processes and influence quality.

HCA indicated that DNA methylation is influenced more by postharvest handling than fruit ripening stage. Fruit quality in contrast, correlated strongly with ripening. The exception was ‘5T’ fruit, where there was the distinct DNA methylation state and quality characteristics implied a strong regulatory mechanism between chilling, ripening and methylation. Broader analyses of the methylome and transcriptome by whole genome bisulfite-sequencing and RNA-Seq (Al Harrasi et al., 2017; Wang et al., 2009), may provide a more comprehensive picture of how early-harvest and low-temperature storage influence tomato ripening at the molecular level.

We proposed a model connecting postharvest strategies and its induced changes in fruit ripening (Fig. 8). Postharvest handling modulates *SlDML2* expression, which in turn influences fruit global DNA methylation. Changes in methylation status may have consequences for a subset of ripening genes (Lang et al., 2017), even if *RIN* expression remained robust in ripened fruit regardless of storage treatment (Fig. 6B). *RIN* is therefore not a reliable proxy for informing on the endogenous or physiological conditions that influence ripening, only that the stage was attained.

**Fig. 8.**
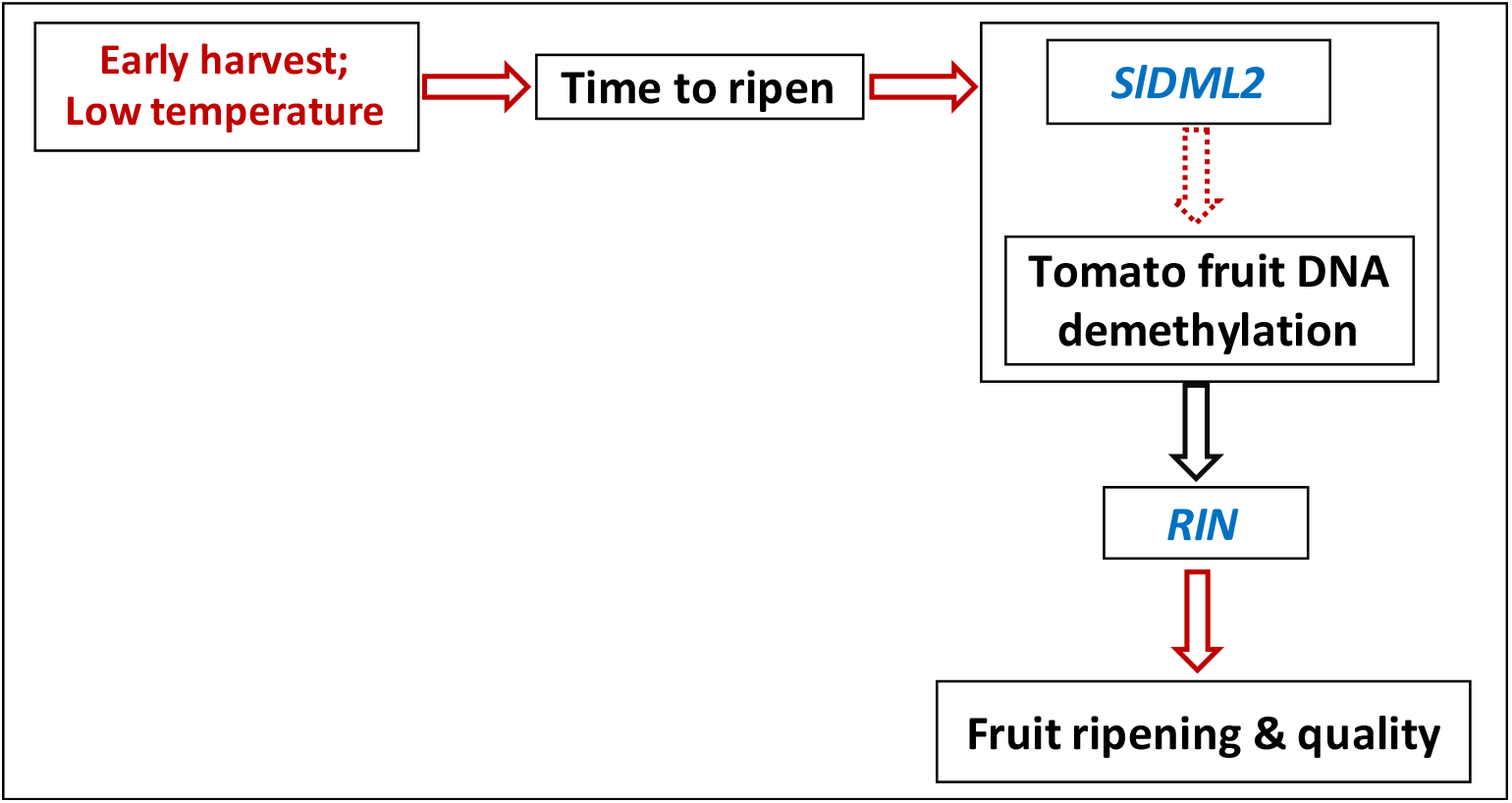
A proposed model showing the relationship between postharvest handling and changes in the tomato fruit development. This model is based on variables (shown as boxes) studied in this work and how they are related. The red arrows indicate positive relationships between variables, and the black arrow indicates no change based on this work. Genes and postharvest conditions are shown in blue and red font respectively. Early harvest and low-temperature storage changed ripening time, resulting in differences in the transcriptional levels of *SlDML2* and changes in global DNA methylation status. However, the relationship between *SlDML2* expression and DNA methylation was not linear (dashed arrow). *RIN* expression remained the same regardless of the different postharvest handling and DNA methylation levels verified in this work, but was related to the ripening stage attained by the fruit. Our postharvest practice widened the fruit ripening/developmental window which influenced *SlDML2* expression.

There are many strategies for prolonging the shelf-life of tomato fruit, but they often reduce flavor. The postharvest treatment used in this study negatively influenced fruit sensory attributes and this may be mediated in part through DNA methylation. In support of this, Zhang et al., (2020) recently found that exogenous ethylene stimulated *SlDML2* transcripts and DNA demethylation. There are also additional epigenetic regulatory mechanisms that may indirectly influence tomato fruit ripening and quality, and it would be of interest to determine how they are affected by postharvest methods (Zhou et al., 2019; Lü et al., 2018; Liang et al., 2020). Greater efforts are needed to help to unravel the complex regulatory ripening network in tomato at the epigenetic and transcriptional level. For example, it will be important to explore the effect of anthropogenic postharvest environments on the timing and dynamics of DNA demethylases and DNA methylation. Such studies may provide novel ways to extend fruit shelf life as well as reduce postharvest loss.

## 5. Conclusion

Our aim was to understand potential changes in DNA methylation and tomato fruit quality in relation to postharvest handling. We have demonstrated that early-harvest and low-temperature conditions significantly reduced fruit quality such as color, the sugar-to-acid ratio and firmness. Expression of the *SlDML2* gene is essential for DNA methylation and fruit ripening, here, we showed that its expression was also responsive to postharvest handling. There were large changes in DNA methylation due to low temperature and early harvest, since the DNA methylation atlas of fruit at the same ripening stage, but developed under different handling regimes, was varied. The relationship between DNA methylation and fruit quality was not linear, and there are likely to be complex biological mechanisms influenced by DNA methylation that control fruit quality. To sum up, postharvest handling induced loss of fruit quality went along with the alterations in global DNA methylation state i.e. demethylation and methylation events.

## Author Credit

**Jiaqi Zhou** - Data curation; Formal analysis; Funding acquisition; Investigation; Methodology; Validation; Visualization; Writing - original draft, review & editing; **Bixuan Chen** - Formal analysis; Funding acquisition; Investigation; Methodology; Writing - review & editing; **Karin Albornoz** - Methodology; Writing - review & editing; **Diane Beckles** - Conceptualization; Funding acquisition; Methodology; Project administration; Resources; Supervision; Writing - review & editing original and final drafts.

## Acknowledgements

We thank Jason Romero, Keqing Wang, and Yiran Wang for assistance in performing the experiments.

