## Supplementary material for "Postharvest handling induces changes in fruit DNA methylation status and is associated with alterations in fruit quality in tomato (*Solanum lycopersicum* L.)": Fig. S1, Fig. S2, Fig. S3, Fig. S4, Table S1, Table S2, Table S3, Table S4

[š] Current Address: United Genetics Seeds, Hollister CA

**List of Figures and Tables**

Fig S1. Fruit developmental stages of cv. Micro Tom

Fig S2. *SlDML2* relative expression measured by RT-qPCR

Fig S3. *SlDML2* expression data of cv. Ailsa Craig from FruitENCODE database

Fig S4. *RIN* expression data of cv. Ailsa Craig from FruitENCODE database

Table S1 Primers in Semiquantitative RT-PCR

Table S2 Primers in quantitative real-time PCR

Table S3 Adaptors and primers used in MSAP

Table S4 MSAP site types and methylation status

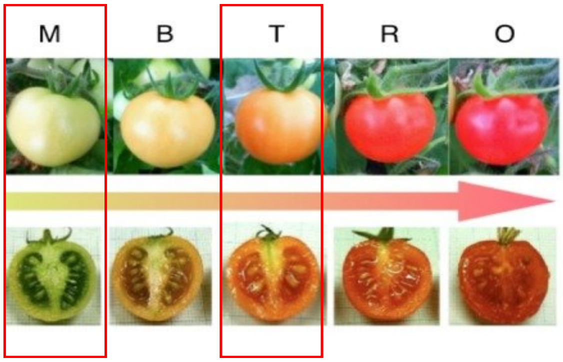


**Fig S1. Fruit developmental stages of cv. Micro Tom (adapted from Takizawa et al., 2014).** Mature Green (M); Breaker (B); Turning (T); Ripen(R) and Over Ripe (O). Mature Green and Turning were tested in this work.

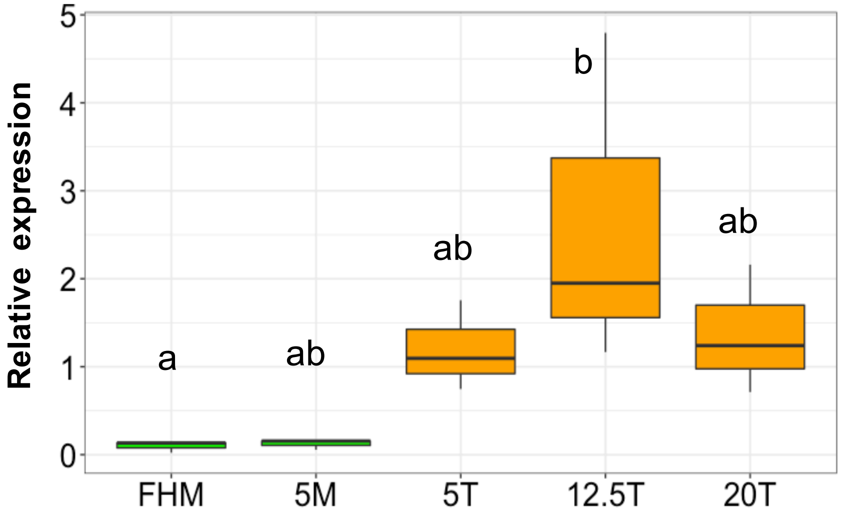


**Fig S2. *SlDML2* relative expression measured by RT-qPCR.** $2^{\Delta\Delta C_T}$ analysis was used, in which *SlAct7* as reference gene and fresh-harvested Turning fruit (‘FHT’) were used as the calibrator. Each treatment includes three biological replicates. Letters above the box indicate significant differences across all groups ($p < 0.05$, CLD).

| Expression (RPKM) | 7 DPA | 17DPA | 27DPA | 37DPA | 42DPA | 47DPA | Leaf |
|---|---|---|---|---|---|---|---|
| Colorless non-ripening (Cnr) | 43.88 | 45.48 | 41.74 | 49.06 | 78.54 | 51.12 | NA |
| green flesh (gf) | NA | NA | 20.54 | 64.86 | 187.28 | 188.23 | NA |
| high pigment (hp) | NA | 16.57 | 29.06 | 48.26 | 178.43 | 104.97 | NA |
| non-ripening (nor) | 26.59 | 20.35 | 28.19 | 19.53 | 23.5 | 28.8 | NA |
| never-ripe (nr) | 27.45 | 21.9 | 30.96 | 48.37 | 242.26 | 81.44 | 10.14 |
| ripening inhibitor (rin) | 36.7 | 31.81 | 31.1 | 52.47 | 95.09 | 122.06 | NA |
| wild-type (Ailsa Craig) | 30.08 | 29.53 | 11.48 | 31.69 | 52.44 | 39.13 | 15.35 |

Mature green Breaker Light red to red ripe

**Fig S3. *SlDML2* expression data of cv. Ailsa Craig from FruitENCODE database** (Lü et al., 2018)**.**

| Expression (RPKM) | 7 DPA | 17DPA | 27DPA | 37DPA | 42DPA | 47DPA | Leaf |
|---|---|---|---|---|---|---|---|
| Colorless non-ripening (Cnr) | 0.0 | 15.35 | 0.0 | 15.45 | 210.79 | 258.03 | NA |
| green flesh (gf) | NA | NA | 0.0 | 11.06 | 157.65 | 576.96 | NA |
| high pigment (hp) | NA | 0.0 | 5.4 | 21.11 | 605.74 | 1173.69 | NA |
| non-ripening (nor) | 0.0 | 0.0 | 2.99 | 0.0 | 13.26 | 14.68 | NA |
| never-ripe (nr) | 0.0 | 0.0 | 2.76 | 17.26 | 614.62 | 338.6 | 0.0 |
| ripening inhibitor (rin) | 0.0 | 0.0 | 1.87 | 5.35 | 232.43 | 663.65 | NA |
| wild-type (Ailsa Craig) | 0.0 | 0.0 | 0.0 | 70.5 | 613.13 | 614.27 | 0.0 |

Mature green Breaker Light red to red ripe

**Fig S4. *RIN* expression data of cv. Ailsa Craig from FruitENCODE database** (Lü et al., 2018)**.**

**Table S1 Primers in Semiquantitative RT-PCR**

| Gene ID | Name | Nucleotide sequence (5'-3') | Product length (bp) |
|---|---|---|---|
| Solyc03g078400.2 | SlACT7-F | GCTATCCAGGCTGTGCTTTC | 157 |
| | SlACT7-R | CAGTAAGGTCACGACCAGCA | |
| Solyc05g012020 | RIN-F | ATTGGGCACAAAAGACTTGG | 212 |
| | RIN-R | CACTTTGCTCACCACAATGC | |
| Solyc10g083630 | DML2-F | ATACAGGCCGTCAACTTTGG | 445 |
| | DML2-R | CCCTTTGGCATTTATGCTGT | |

**Table S2 Primers in quantitative real-time PCR**

| Gene ID | Name | Nucleotide sequence (5'-3') | Product length (bp) |
|---|---|---|---|
| Solyc03g078400.2 | SlACT7-F | GCTATCCAGGCTGTGCTTTC | 157 |
| | SlACT7-R | CAGTAAGGTCACGACCAGCA | |
| Solyc10g083630 | DML2-F | GCAGCAGTTCATGCTTACCA | 95 |
| | DML2-R | CCCTTTGGCATTTATGCTGT | |

**Table S3 Adaptors and primers used in MSAP**

| Name | Nucleotide sequence (5'-3') |
|---|---|
| *EcoR*I adaptors | CTCGTAGACTGCGTACC<br>AATTGGTACGCAGTCTAC |
| *Hpa*II/*Msp*I adaptors | GATCATGAGTCCTGCT<br>CGAGCAGGACTCA TGA |
| *EcoR*I Preselective primer | GACTGCGTACCAATTC |
| *Hpa*II/*Msp*I Preselective primer | ATCATGAGTCCTGC TCGG |
| Selective primer pair 1 | GACTGCGTACCAATTCACC<br>ATCATGAGTCCTGCTCGGTCAA |
| Selective primer pair 2 | GACTGCGTACCAATTCACC<br>ATCATGAGTCCTGCTCGGTCCA |

**Table S4 MSAP site types and methylation status**

| **Site type** | ***Msp*I** | ***Hpa*II** | **MSAP Site Type** | **Methylation State** |
|---|---|---|---|---|
| I | 1 | 1 | Demethylated | CCGG<br>GGCC |
| II | 0 | 1 | Hemi-methylated external C | **C**CGG<br>GGCC |
| | | | Hemi-methylated both C | **CC**GG<br>GGCC |
| III | 1 | 0 | Fully methylated internal C | C**C**GG<br>GG**C**C |
| IV | 0 | 0 | Fully methylated both C | **CC**GG<br>GG**CC** |
| | | | Fully methylated external C | **C**CGG<br>GGC**C** |

| | | | | |
|---|---|---|---|---|
| | | | Fully methylated external C +<br>Hemi-methylated internal C | **CC**GG<br>GGC**C** |
| | | | Hemi-methylated external C +<br>Fully methylated internal C | **CC**GG<br>GG**C**C |
